# Environmental factors and cultural drift influence song evolution in New World Sparrows

**DOI:** 10.1101/2023.09.01.555954

**Authors:** Kaiya L. Provost, Jiaying Yang, Bryan C. Carstens

**Affiliations:** Department of Biology, Adelphi University; One South Avenue, Garden City NY, 11530, USA; Museum of Biological Diversity and Department of Evolution, Ecology and Organismal Biology, The Ohio State University; 1315 Kinnear Rd., Columbus OH, 43212, USA; Biological Science Department, Vanderbilt University; 2201 West End Ave, Nashville, TN 37235, USA

## Abstract

Variation in bird song is often assumed to be determined by sexual selection, rather than natural selection. However, most investigations to date have drawn their conclusions from a handful of species due to the challenges with manually processing sound data. Here, we use deep machine learning to investigate nearly all species of New World Sparrows. We leverage existing data to identify the processes that structure variation in bird song and to determine how this variation corresponds to patterns in genes and traits. Song variation in ~40% of species can be explained by environment, geography, and time. Across a community and global scale, the action of natural selection on the evolution of song is at least as impactful as it is on other genetically-determined traits.

## Main Text

Singing behavior in birds has a high impact on reproduction, fitness, and speciation, especially in Passeriformes (*1–3*), but it remains unclear how song evolves within species. Hypotheses generally explain the evolution of bird song via the action of selective or neutral forces. For example, natural, sexual, and social selection have been invoked by many researchers (*4–6*), including adaptation to the environment (*7–10*). Stochastic evolution, either via cultural drift or song learning, has also been reported in multiple investigations (*11–15*) with genetic drift implicated in others (*16–22*). Similar to the selectionist / neutralist debates in molecular evolution (e.g., *23*) this question has been difficult to disentangle in large part due to the small datasets that have been applied to test these hypotheses. Most researchers rely on recordings of bird songs collected from the field, converted to audiograms, and manually processed to measure differences in the timing, placement, and frequency of the syllables that constitute the songs (e.g., Fig. 1). However, audiogram data is laborious to collect, with each syllable taking on average four seconds to annotate based on preliminary work.

**Fig. 1:**
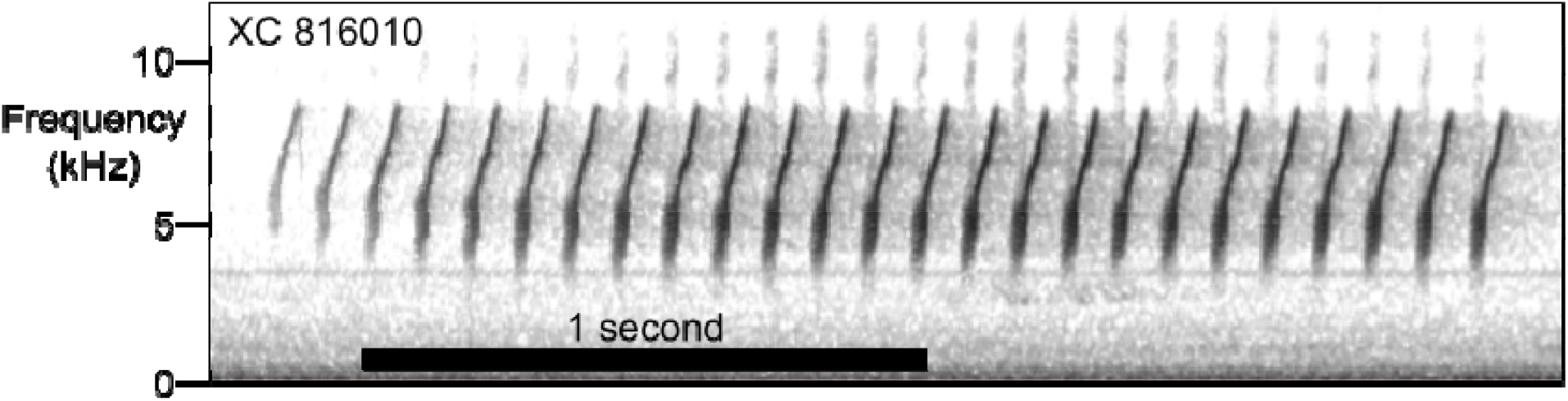
Example spectrogram from Chipping Sparrow (*Spizella passerina*). Recording depicts the first syllable of XC 816010 with 25 individual notes. X-axis depicts time in seconds, with black bar showing one second of time. The Y-axis depicts frequency (kHz). Darker colors indicate sounds are louder – note that gray color below the notes represents background noise, while gray color above notes represents harmonics.

We test what factors predict intraspecific diversity in songs by quantifying intraspecific variation and its correlates in the New World Sparrows (Family: Passerellidae). This family was chosen because they reside in a variety of habitats and are notable for their variation in song across species. They are also well-studied in our existing curated song datasets (e.g., Xeno-Canto, Borror Lab of Bioacoustics, Macaulay Library). For example, on Xeno-Canto at the time of writing, the 16,833 Passerellidae recordings make up 2% of the total ~842,000 bird recordings, despite comprising only 1.2% (134) of the 10,495 bird species. The Borror Lab of Bioacoustics database has an even higher bias: 31.8% (14,360/45,097) of the bird songs are Passerellids. Data on intraspecific variation in bird songs can be challenging to extract from the mountain of preexisting recording data. We labeled ~500,000 individual song syllables across ~26,000 recordings and 125 species (*24, 25*). We calculated song properties for each syllable to estimate song variation. Properties include frequency, frequency bandwidth, number of inflection points within a syllable, slope of the syllable, time duration, etc. These properties can differentiate syllables of different shapes. Rather than devoting ~24 days to process ~500,000 syllables, we applied a deep learning approach to automate the extraction of data from these recordings.

Convolutional neural networks (CNNs) are a type of deep machine learning algorithm that take image data and summarize across subsets of the images to identify important features in the image (see *26, 27*). CNNs and other deep learning algorithms can be trained to automate auditory data collection because sound is easily converted to images with little loss of information (*28*). Different sounds therefore have different shapes on a spectrogram (Fig. 1). The CNN can learn the shapes we are interested in, then automatically identify those shapes. The corresponding increase in the amount of data that can be analyzed is comparable to that which was enabled in molecular genetics by advances in DNA sequencing technology that enabled researchers to move from single locus data to whole genomes.

### Heterogenous factors determine song variation within species

In 61% of New World Sparrows, song was not associated with the environment, year of recording, or distance between recordings (Fig. 2). For the remaining 39%, the factors determining variation in song within species were highly heterogeneous across Passerellidae. Song was significantly associated with the environment in 19% of species, year of recording in 17% of species, and distance between recordings in 14% of species, with 12 species showing multiple significant factors. Significant associations were more likely to be detected in species with higher absolute song differentiation within-species (p=0.031). Genetic differentiation is known to be associated with song differentiation (*15, 16*). We performed a secondary analysis on a subset of 28 taxa with intraspecific genetic data using phylogatR (*29*). For 71% of species, intraspecific genetic variation is not associated with environment or geographic distance. For the remaining 29%, half of the species showed associations with the environment and half showed association with geographic distance. For the 28 species that had both song and genetics evaluated, 43% showed no associations between intra-specific variation and environment or distance between recording. 29% had associations between song but not genetics, and 21% associations between genetics but not song. For the remaining species, song was associated with geographic distance but genetics was associated with environment. There is evidence that genetic distance and song distance in other species is correlated (e.g., *30*), but in our dataset it seems that generally they are not being caused by the same factors.

**Fig. 2:**
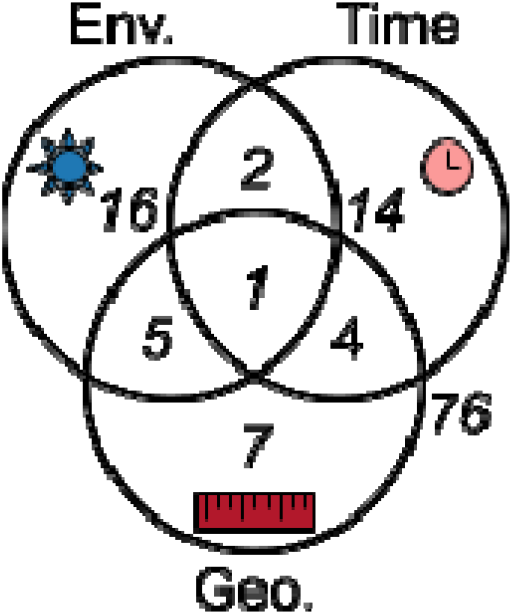
Venn diagram displaying proportion of species whose songs are best predicted by different features. Features are Environment (“Env.” on plot, blue sun), Time (pink clock), and Geography (“Geo.” on plot, red ruler). Number outside of circles indicates number of species with no significant association between song and features.

Using existing morphological data (*31*), we find that species whose songs are associated with the environment have longer wings, longer tails, and longer secondary feathers (p<0.0291), as well as being generally larger but not significantly so (p=0.056). Flight feather morphology is often used as a proxy for dispersal in birds (*32*), with longer and pointier wings indicating better presumed dispersal ability. This suggests that the influence of spatial factors on diversification within species (i.e., in songs or genetics) is particularly pronounced on birds with poorer dispersal ability.

### Explanations of song variation are not associated with phylogeny

No major patterns emerge from an examination of phylogenetic relationships within the context of the spatial analyses conducted here. While song production itself has a phylogenetic component (*33*), the impact of phylogeny on the specific attributes of bird song is unclear. Further, there is evidence that divergence at different taxonomic levels is not due to the same process (e.g., *34*). If phylogeny is a large driver of spatiotemporal features in bird songs, we would expect to find clades where one spatiotemporal feature was prominent in its impact. While we see this within some genera (e.g., Atlapetes has a high proportion of taxa whose song is associated with the environment), such a pattern does not appear at larger scales (Fig. 3). Our results are thus consistent with phylogeny having some small effect on the evolution of bird song, but additional work is needed to explore higher level effects in comparative phylogenetic contexts. One note is that taxonomic uncertainty is a challenge to studies like the one conducted here. As a potential hedge against the analysis of cryptic species, we performed the same analyses using subspecies as our focal taxa (see supplement) and found similar results, though a smaller proportion of taxa have significant relationships between spatiotemporal features and song likely due to sample size. That the patterns are similar suggests that our findings are robust to taxonomic uncertainty.

**Fig. 3:**
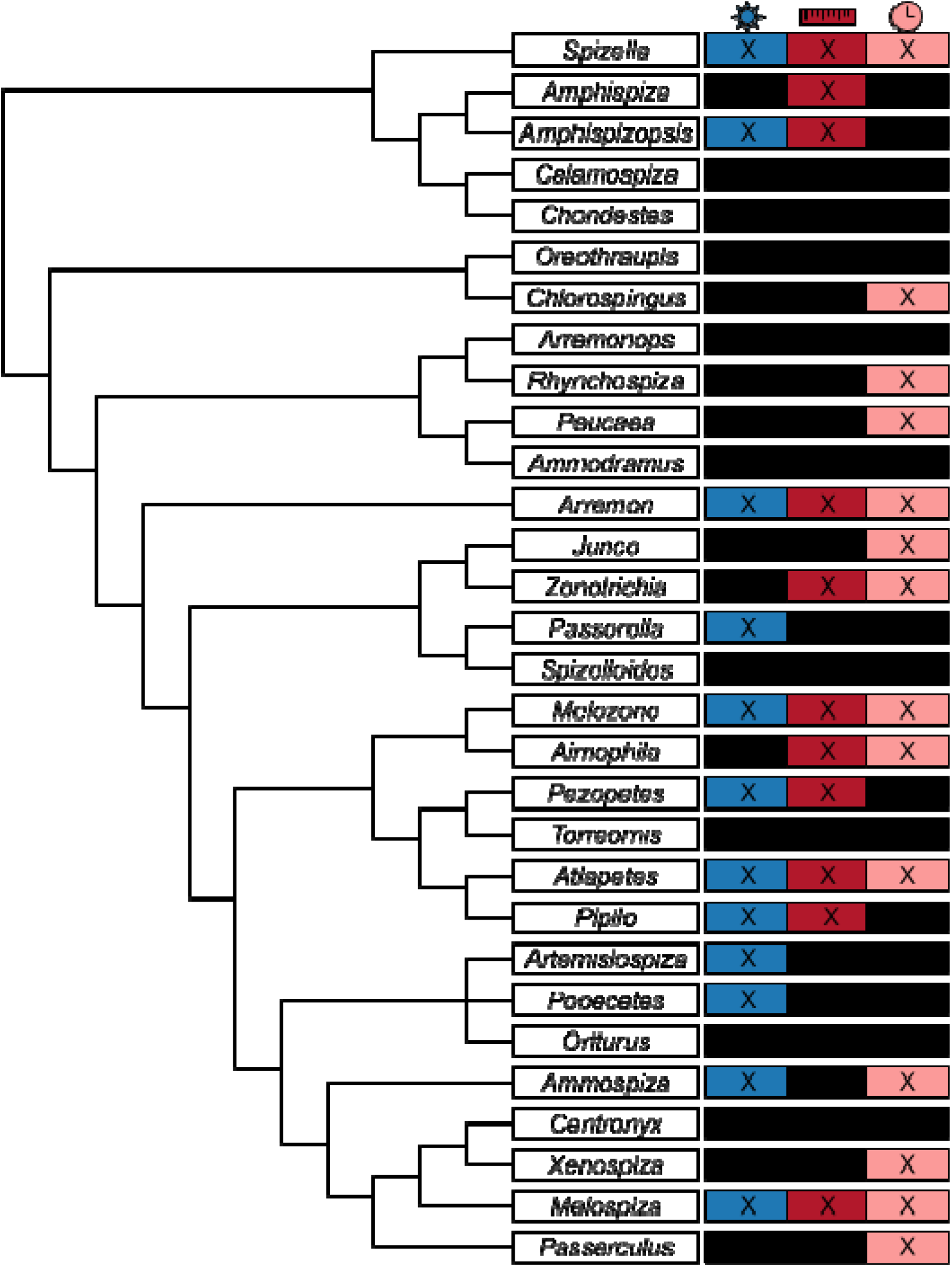
Best predictors of species-level differences in traits. Left: phylogeny at the generic level, with sample sizes of species sampled/total species. Right: heat map of number of species sampled that show significant differences in environment (sun, left, blue), geographic distance (ruler, center, red), or time (clock, right, pink). Colored values with an “X” indicate at least one species shows significant differences, black values indicate no species shows significant differences.

For the taxa wherein neither of these factors is important in explaining song variation, there is still work to be done to determine what is influencing song evolution. Across many species, there appears to be little song variation. This could result from small sample size, but also could be an indication that song variation is not present. In contrast to taxa with little song variation, there are a few taxa that have high song variation that are not explained by stochastic or environmental factors (e.g., *Pipilo maculatus*). One explanation might be that these species do not occupy environmentally variable areas, perhaps due to very strict niche requirements. However, we don’t find any evidence that absolute environmental differences are driving these patterns. One further avenue to explore is that of acoustic niche partitioning (*35*), wherein species occupying the same areas differentiate their songs so as to minimize interference with other species’ signals. This is important in both tropical bird communities (e.g., *36, 37*) and temperate ones (e.g., *38*).

Lastly, it should be noted that our data are unable to investigate sexual and social selection, which are traditionally thought to be important determinants of song evolution across Passerines and birds more broadly (*4*).

Automatic processing of bioacoustic data at the family level has allowed us to address questions related to the factors that influence song evolution across a phylogenetic scale. The results presented here are, to our knowledge, the first large-scale intra-specific analysis of an entire bird family. Overall we find that intraspecific variation in song is not broadly correlated with any particular factor we tested, as the majority of Passerellidae species don’t show correlations with environment, time, or distance. In species where there are correlations, it appears to be driven by different factors and is idiosyncratic across the phylogeny. Though it is possible that these results are dependent on data sampling, particularly since we are using repurposed data, our results suggest that the process of bird song evolution is not a deterministic one. Intraspecific diversity in songs is not explained entirely by phylogeny, geography, cultural drift, or environment. Rather, these factors contribute such that each individual taxon has its own unique response.

## Acknowledgments

We thank E. Akçay, R. Duckworth, and one anonymous reviewer for feedback on an earlier version of this manuscript. Resources from phylogatR and Imageomics were made possible by funding from the NSF and the Imageomics Institute. We thank all individuals who have contributed to song repositories over the last century, especially those at the Borror Laboratory of Bioacoustics.

## Funding

National Science Foundation DEB-1910623 (BCC)

National Science Foundation DEB-2016189 (BCC, KLP)

National Science Foundation OAC-2118240 (BCC)

## Author contributions

Conceptualization: BCC KLP

Data curation: KLP

Formal analysis: JY KLP

Funding acquisition: BCC

Investigation: KLP

Methodology: JY KLP

Resources: BCC KLP

Visualization: JY KLP

Writing – original draft: KLP

Writing – review and editing: BCC JY KLP

## Competing interests

Authors declare that they have no competing interests.

## Data and materials availability

Datasets will be available upon acceptance. Code is available at https://github.com/kaiyaprovost/bioacoustics/

## Supplementary Materials

Materials and Methods

Supplementary Text

Figs. S1 to S8

Tables S1 to S3

## Notes

### Competing Interest Statement

The authors have declared no competing interest.

### Summary of Updates

Revision after peer review and rejection

