## Supplemental Text for "Environmental factors and cultural drift influence song evolution in New World Sparrows"

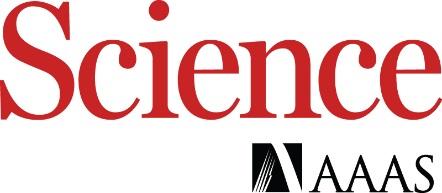

Supplementary Materials for

**Title: Environmental factors and cultural drift influence song evolution in New World Sparrows**

Kaiya L. Provost, Jiaying Yang, Bryan C. Carstens

**The PDF file includes:**

Materials and Methods

Supplementary Text

Figs. S1 to S8

Tables S1 to S3

### Materials and Methods

#### Focal System: Passerellidae

We focused on the New World Sparrows, Family Passerellidae. Our work adopted the IOC taxonomy for determining species designations (*39*). Under this framework, the New World Sparrows include 30 genera, 138 species, and 419 named subspecies (*39-42*). The clade is distributed throughout the entirety of North and South America, save for the northernmost latitudes, and occupying a broad range of habitats therein (*43*).

Phylogenetic relationships within this group vary, though taxa generally fall into eight major clades (i.e., *40*). We used one recent phylogeny (*42*) at the genus level to visualize phylogeographic patterns. Since some genera were missing from that phylogeny, the placement of those genera followed two previous phylogenies (*41, 40*). The genus *Xenospiza* was placed as sister to *Centronyx*, which was in turn sister to *Melospiza*. The genus *Torreornis* was placed as sister to *Pezopetes*, and similarly *Oreothraupis* as sister to *Chlorospingus* and *Amphispizopsis* as sister to *Amphispiza*.

#### Acoustics Data Processing

We downloaded all 10,836 songs of the family Passerellidae from Xeno-Canto (XC) on 5 July 2022. We supplemented these recordings with 12,145 songs from the Borror Lab of Bioacoustics (BLB) collection. In both cases, previous studies had annotated songs from these repositories (*25, 44*); we used those annotations to supplement our analysis. The few individuals who were described in the metadata as hybrids were eliminated from the analysis. Songs were converted to mono WAV format and to a sample rate of 48,000 Hz using the resamp function in the seewave 2.2.0 package in R version 4.2.1 (*45, 46*).

Recording quality varied across taxa, with some being of high quality (e.g., Xeno-Canto quality A), low quality (e.g., Xeno-Canto quality E) or unknown quality (e.g., the BLB recordings). We tested the impact of recording quality by repeating MMRR analyses (see below) on subsets of homogenous recording quality at the family level.

#### Machine Learning

We used a machine learning model (i.e., TweetyNet; *24*) to identify the locations of individual syllables in each recording. A model that had been previously trained on nine species, including non-Passerellids (i.e., 9SppBalanced; *25*) was applied to the new analysis of recordings three ways. First, the model was used to identify syllables without making any changes to the model. Second, we fine-tuned the model on only Passerellidae species with annotations from another study (*25*) and used that version to identify syllables. Lastly, we fine-tuned the model again on all annotated Passerellidae that we had generated as part of the present study. After each set of predictions of syllables had been made, we extracted as syllables anything that the model had identified across any of the three sets of predictions. We removed all syllables that were less than 0.05 seconds in length as spurious.

#### Summary Statistics

In a similar manner to how phenotypic traits are measured within samples before being included in a quantitative analysis, we extracted summary data from the song syllables. This was accomplished using the warbleR package version 1.1.28 (*47*) in R version 4.2.1. We calculated summary statistics based off of spectrograms using the function spectro_analysis. The summary statistics included metrics of song duration; mean and standard deviation of frequency; quantiles (including median, 25%, and 75%, and the interquartile distance) of frequency; quantiles of time duration; skew and kurtosis of the frequency spectrum; entropy measurements (spectral entropy, time entropy, and overall entropy); spectral flatness; dominant frequency calculations (mean, minimum, maximum, range, and value at start and end of sound); modulation index and slope of dominant frequency; peak frequency; and mean peak frequency. We also calculated fundamental frequency time series using the function freq_ts, which allowed us to get the frequency at each time bin. This lets us calculate the number of inflections, slopes, and mean slope per sound.

After generating these summary statistics, we extracted five for further analysis: bandwidth, as the IQR of frequency; time duration, as the IQR of the time; center, as the median frequency; inflections, as the number of fundamental frequency inflection points; and slope, as the mean fundamental frequency slope. From these five summary statistics we calculated values for each syllable, then generated a centroid value for each individual across all syllables. We performed a principal components analysis (PCA) on these data for downstream analyses.

Once centroid values for the five summary statistics were calculated for all individuals, we calculated euclidean distances between all individuals across all species. These euclidean distances were used as representation of song variation as the predictor variable of the song MMRR (see below). We then created subsets of these euclidean distances for each genus, species, and subspecies across Passerellidae. Finally, to serve as a proxy for the overall distances represented within each taxon, we took the mean and standard deviation of euclidean distance for each.

#### Genetics

We downloaded geospatially labeled genetic data for all birds from GenBank using the phylogatR framework (*29*) on 7th December 2020. Of the 9,338,693 sequences present for all birds, Passerellidae make up 29,030 of those sequences representing 36 species. The overwhelming majority of the genes were CO1, so we restricted our analyses to that gene only.

From these genetic data we calculated genetic distances between individuals using the dist.gene function in the ape package version 5.7 in R (*48*). After that, we used those genetic distances as the predictor variable in our genetics MMRR. We also calculated mean genetic distance within species by taking the average pairwise across all individuals. We chose to do genetic analyses as a parallel because the song data and genetic data were not always collected from the same localities and as such cannot be directly compared in the same model. Instead we chose to compare and contrast the two models to see if similar patterns emerge.

#### Spatiotemporal

To examine the impact of the environment, we downloaded rasterized data from multiple sources representing various environmental features; climate, land cover, habitat type, topography, soil features, urbanization, geographic distance, and temporal distance (*49-54*). After removing variables that were invariable in our dataset, this resulted in 48 total variables. For all song and genetic data points with latitudes and longitudes, we extracted the environmental variables for each point. We performed a PCA on the environmental data, centering and scaling the variables and imputing missing data such that any missing values were assigned the mean. We did this using the prcomp function in the stats package in R. We chose to retain five of the resulting PCs as they explained over 50% of the variation across the 48 original variables, and these were used to represent environmental distance. We then calculated a euclidean distance matrix of environmental distance between all individuals in both the genetic and song datasets.

We also calculated pure geographic euclidean distance between individuals from the latitude and longitude data, which serves as a metric of geographic distance in the MMRRs. While most longitudes were highly negative, some on the very western tip of Alaska were highly positive due to the transition from 180°W longitude to 180°E longitude, artificially inflating their geographic distance. To correct for this, individuals in this region had their distances corrected. For example, an individual with 172.0°E longitude would be converted to 188°W longitude, or -188 decimal degrees longitude. We treated geographic distance as representing the stochastic process of isolation-by-distance.

Finally, we also calculated a temporal distance between individuals based on the year a recording was made. Note that year of recording was only available for the song data: we did not have information for our sequences as to when samples were collected. We treated temporal distance as representing the stochastic process of cultural drift.

#### Synthesis

We examined biases in sampling in XC, BLB, and GenBank data for Passerellidae using the locational and temporal metadata associated with each.

We performed a multiple matrix regression with randomization (MMRR) analysis to test our hypotheses (*55*). MMRR estimates the coefficients of relationships between multiple distance matrices, controlling for the impact of all of the variables in the dataset, and then uses a randomization technique to estimate a *p*-value of significance. When an MMRR analysis is conducted with the same input matrices, all of the returned coefficients are identical except for *p*-values. We conducted MMRR comparisons multiple times for each dataset, randomizing each 1000 times, and then averaged the *p*-values accordingly. For the song models each were set up so that song distance was being predicted by environmental distance, geographic distance, and temporal distance. Because we did not have temporal information for our genetic data, those models were set up so that genetic distance was predicted by environmental and geographic distance only. If the environmental selection hypothesis is supported, the MMRR will return a significant value *p*-value (using an alpha value of 0.05) for the environmental distance matrix. If the stochastic cultural drift hypothesis is supported, the MMRR will return a significant p-value either geographic distance, temporal distance, or both. Both hypotheses may also be supported simultaneously.

MMRRs were performed at four levels: family, genus, species, and subspecies. We performed these analyses at these different taxonomic levels to evaluate the impact of each factor on progressively larger populations. We then compared taxa that were significant in environment or geography for both song and genetics with their morphological measurements. We also compared this significance with the absolute differences present in genetics and song across species. We look at the phylogenetic relationships as well to see whether there is evidence that these patterns are phylogenetically conserved. We also tested the impact of recording quality by repeating MMRR analyses on subsets of homogenous recording quality.

For the full-family dataset, we were unable to run our MMRR models on the full distance matrices as they were too large. Instead, we chose to calculate distance matrices using genus, species, and subspecies average, as well as for only individuals at each of the five song quality types that XC provided (from high-quality “A” to low-quality “E”). This resulted in eight datasets to evaluate the impacts at the family-level.

#### Morphology

Morphological differentiation, and phenotypic differentiation more broadly, is one major way that species adapt to their environment. In birds, bill morphology is thought to be selected with respect to diet (*56, 57*), song (*58*), nest building (*59*), and thermoregulation (*60, 61*). Bill morphology is also constrained by phylogeny (*62*). Like in other vertebrates, body size is a key factor for thermoregulation (*63, 64*), and is also important in influencing song (e.g., *65*). To assess the impact of morphological measures on our data, we downloaded data from AVONET (*31*), which has measures of beak length (to nares and culmen), beak width, beak depth, tarsus length as a proxy for body size, primary and secondary feather length, tail length, the hand-wing index, and Kipp’s distance.

This dataset contained 127 Passerellidae species across all 30 genera (missing: *Arremon basilicus, A. costaricensis, A. perijanus, A. phaeoplerus, Atlapetes meridae, At. nigrifrons, Junco bairdi, Passerella megarhyncha, P. unalaschcensis,* and *Pipilo naufragus*). There were 1,177 measurements taken across the family. We used the average value for each species as a factor when looking at the genetic and song MMRR results. We also calculated the average value for each genus. We evaluated whether morphology explained our results by using ANOVA and Tukey’s Honest Significant Difference tests (*66, 67*). We evaluated significance using an alpha value of 0.05.

### Supplementary Text

#### Focal System: Passerellidae

Sampling of the Passerellidae is uneven across North and South America in both song and genetics (Supplemental Figure 1). The western coast of North America, and Ohio, are highly sampled, whereas much of the Amazon, central North America, and the Caribbean are undersampled. The recordings vary in terms of time recorded as well (Supplemental Figure 2). The BLB is a historical collection and substantially older recordings (range 1948-2014, mean 1979) than XC (range 1976-2022, mean 2013). Both databases also have peak recording from April to July, which corresponds to the breeding season for most species. 80.0% of BLB songs were recorded during these months, whereas 56.3% of XC songs were recorded during these months. Across space, sampling between the recording data and the genetic data is significantly correlated though the correlation is weak (log-corrected, p=0.030, adjusted R^2^ = 0.038).

#### Acoustics Data Processing

After processing all song recordings through the TweetyNet model (*24*), our data consisted of 26,169 recordings and 513,617 annotated syllables. This comprised all 30 genera, 134 species, and 316 subspecies. All but one recording had latitude and longitude information, and 23,544 had date information. Taxa ranged in terms of their overall song differentiation. At the species level, the range was from 0.00 (*Chlorospingus inornatus*) to 4.32 (*Atlapetes personatus*). At the genus level, mean song distances ranged from 1.64 (*Passerella*) to 3.37 (*Ammodramus*). At the subspecies level, the range was again from 0 (*Atlapetes nationi brunneiceps, A. semirufus benedettii, Melozone crissalis senicula, Pipilo maculatus curtatus*) to 5.02 (*Peucaea ruficauda lawrencii*).

#### Summary Statistics

We extracted qualitative metrics from our dataset. For our PCA on song variables, we retained all five PCs for downstream analyses (Table S1). Syllables with high values of PC1 were long, with numerous inflection points, and narrow bandwidth. Syllables with high values of PC2 were high-frequency, wide-bandwidth. Syllables with high PC3 had negative slopes and high frequencies. Syllables with high PC4 had narrow bandwidths and positive slopes. Syllables with high PC5 had low frequencies, were short, and had numerous inflection points. We then calculated distances from these data.

#### MMRR Results

For the MMRRs using song data there are 30 genera, 125 species, and 171 subspecies with sufficient information to test the significance of environment, geography, and time (Fig. S3, Table S1, Table S2).

There is a strong correlation with song differentiation (both with and without log-correcting for sample size disparity) such that taxa with more recordings are more likely to have a significant correlation detected (p<4.9x10^-6^).

At the species level, we find that we are more likely to detect significant differences between song and environment when the absolute song differences were higher (p=0.031). Again, this is easily explained, as when there is little song variation whatsoever the likelihood of detecting differentiation is slim. 76/125 species showed no significant relationships between our spatial factors and song differentiation (Main Text Figure 5; Fig. S4). Of the rest, 37/125 had one factor significant: 16/125 were environment, 14/125 were time, and 7/125 were geography. An additional 11/125 had two factors significant: 5/125 for environment and geography, 4/125 for geography and time, and 2/125 for environment and time. The remaining one species had all three significant. This thus resulted in 24/125 species having the environment being significant in some way, 21/125 having time significant, and 17/125 having geography significant.

For our genus-level MMRR analyses, 13/30 genera showed no significant relationship between song and environment, geography, or time (Fig. S1). Ten genera showed one of these factors being significant: 5/30 had time as significant, 4/30 had environment, and 1/30 had geography. An additional four genera had two factors as significant: 2/30 both environment and geography, 1/30 both environment and time, and 1/30 both geography and time. The remaining 3/30 genera had all three factors as being significant. All in all, that resulted in 10/30 of genera with environment being significant in some way, 10/30 with time significant in some way, and 7/30 with geography significant in some way.

We find we are more likely to detect significant associations between song and time when the mean time difference is larger and more variable at the genus level (p<0.0015). This is easily-explained, as when there is little variation in the year of the recording then there are no temporal differences to compare. Song and time are also more likely to be related when geographic distance is higher, but this is not significant (p=0.0584).

At the subspecies level, 132/171 subspecies showed no relationship between song and spatial factors (Fig. S1). Of the remaining, 27/171 had one factor only: 16/171 showed time only, 7/171 showed geography only, and 4/171 showed environment only. Eight subspecies had two factors predicting song: 4/171 had environment and geography, 3/171 had geography and time, and 1/171 had environment and time. The remaining four subspecies had all three significant factors. In total, 13/171 had environment play a role, 18/171 had geography play a role, and 24/171 had time play a role.

At the family level, our results were mixed. When averaging over genera (N=30), we found no significant relationships between songs and environment, geography, or time. However, when averaging over species (N=125), environment was significant, and when averaging over subspecies (N=171) both environment and time were significant.

If we look at subsets of the data corresponding to different data qualities, we find that for high quality songs (N=4,088 “A” and N=3,527 “B”) both environment and geography are significant, but for lower quality songs (N=1,099 “C”, N=225 “D”, and N=62 “E”) nothing was significant.

#### Genetics

After compiling all of the genetic data, we ended up with 428 COI sequences across 20 genera and 36 species. There was no subspecific information available to us. For the genetic MMRRs, there are 15 Passerellidae genera and 28 species with sufficient genetic information to test the significance of stochastic and environmental features (Fig. S4, Table S1, Table S2). Genetic differentiation also ranged across taxa. At the species level, mean genetic distance ranged from 0.00 (*Chlorospingus flavopectus*) to 164.19 (*Arremon taciturnus*). Mean genetic distance for genera ranged from 28.67 (*Aimophila*) to 150.95 (*Arremon*).

Only 8/28 species showed any influence of spatial factors on genetics: four with environment only, four with geography only. For the genera, only 3/15 showed any influence of spatial factors on genetics: one genus had geography predicting genetics, one had environment, and one had both.

We found no correlation between sample size and whether or not genetic differentiation was significant (p=0.751).

#### Spatiotemporal

After performing our PCA of environmental variables, we retained the first five PCs as they explained 53% of the data (Table S2, Table S3). High values of PC1 were associated with regions that were dry, cold, had relatively narrow daily temperature but high annual temperature ranges, and high temperature seasonality. High values of PC2 were associated with regions that were cold and wet, with narrow daily temperature and yearly precipitation ranges, acidic sandy soils, and lots of trees to the exclusion of shrubs, bare cover, grazing land, and irrigated land. High values of PC3 were associated with more urbanized areas, high in croplands and agriculture, with lots of seasonal water, low terrain ruggedness, and shallow slopes. High values of PC4 were associated with cold summers, rough terrain, and lots of permanent water. Lastly, high values of PC5 were associated with high population areas with lots of built up concrete, low elevations, and low relief.

We investigated whether or not absolute differences in our traits and spatiotemporal features influenced the final results. Means and standard deviations are correlated with each other for each variable (range r: 0.72-0.93). Because of this, we only examine significant associations between mean values. Geography and environment are also highly correlated with each other, as expected (r=0.65). All other variables are correlated with each other to lesser extents (r<0.37).

#### Synthesis

We compared whether song and genetics were explained by the same factors for the 28 species that had both genetic and song MMRRs (with the caveat that we did not evaluate genetic influence of time).

For species, there were no taxa that had both gene and song explained by environment. However, 18/28 species had neither gene nor song explained by environment, with the remaining species being mismatched (four species with genes explained by environment, six species with song being explained). Likewise, there were no taxa that had both gene and song explained by geography. 20/28 species had neither gene nor song explained, the remaining eight were mismatched (four with genes being explained by geography, four with song being explained).

For genera, one genus had song and genetics both be explained by environment (*Zonotrichia*); notably this genus found significant influences of both spatial factors for genetics and all three spatial factors on song. Eight genera had song and genetics both not be explained by environment -- of these four were explained by time. Six genera had a mismatch: environment explained one trait but not the other (for one genus it explained genetics, for the other five it explained song; three of the latter were also explained by time).

We now compare differences in geography for genera. One genus (*Zonotrichia* again) had both song and genes explained by geography. Ten genera had neither song nor genes explained by geography -- three of these genera had time explaining song. Four genera were mismatched -- one genus had gene explained by geography but not song, and three genera had song explained by geography but not genes. All four of these genera had time as a factor. 47% of genera having the exact same pattern between genes and song and 40% having a partial match (note that this ignores the temporal aspect present only for song).

#### Morphology

We found that there were some differences in morphology between taxa that showed environmental and stochastic influences on their genetics and song.

At the species level, both vocalization and genetic differences were associated with morphology. In species that showed significant differences in genetics across geographic distances, both hand-wing indices and Kipp’s distances were much shorter (p<0.029), which generally indicates that these birds have rounder, less pointed wings. With respect to vocalization, species that had significant correlations between song distances and environmental distances had longer wings (p=0.012), longer tails (p=0.0291), and longer secondaries (p=0.0275). Body size was also larger, though not significantly so (p=0.056). This is consistent with these birds being larger.

At the genus level, there were few differences across categories, which may be due to examining differences in trait averages. The only major difference is that beak lengths (both to culmen and to nares) were on average shorter in genera that showed significant differences in song across geographic distance (p<0.0495).

#### Phylogeny

We plotted the number of significant associations on the genus-level phylogeny to visualize if there is any bias. Though we do not evaluate this in a statistical framework, we do not detect any phylogenetic patterns in the New World Sparrows at the species, genus, or subspecies levels (Fig. S5, Fig. S6, Fig. S7).

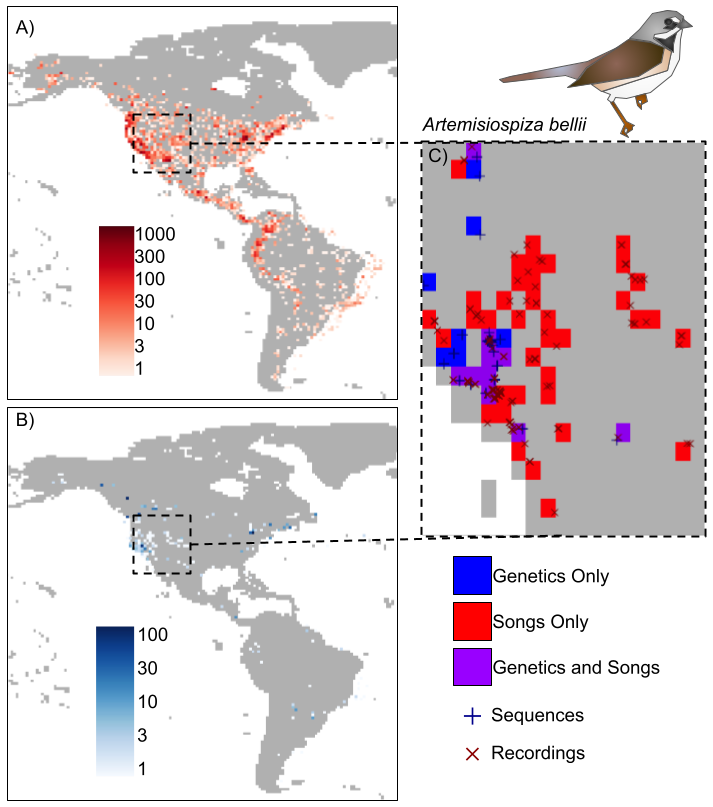

**Fig. S1**.

Song and genetic sampling rarely overlap within species. A) Song sampling from Xeno-Canto of all Passerellidae species. Color indicates density of recordings on a log scale, with lighter reds showing areas with few recordings, darker reds showing areas of many recordings, and gray showing areas with no recordings. Cells are one-degree latitude by one-degree longitude. B) Genetic sampling from phylogatR of all Passerellidae species. Color indicates density of recordings on a log scale, with lighter blues showing areas with few sequences, darker blues showing areas of many sequences, and gray showing areas with no sequences. Cells are one-degree latitude by one-degree longitude. C) Exemplar overlap for one species, *Artemisiospiza bellii*. Points show individual sequences (blue cross) or song recordings (red X). Color beneath shows whether the cell contains genetic data only (blue), song data only (red), or both genetic and sign data (purple). Cells are one-degree latitude by one-degree longitude.

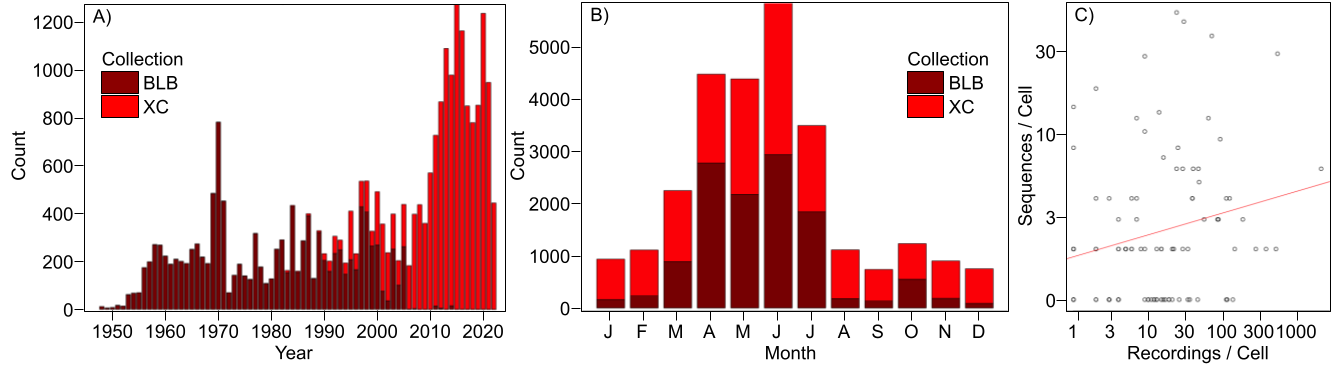

**Fig. S2.**

A) Recordings in the Borror Laboratory of Bioacoustics database (BLB, dark red) and Xeno-Canto (XC, red) across years. B) Recordings in the BLB (dark red) and XC (red) across months of the year. C) Recordings per cell vs sequences per cell across range of Passerellidae, log scaled.

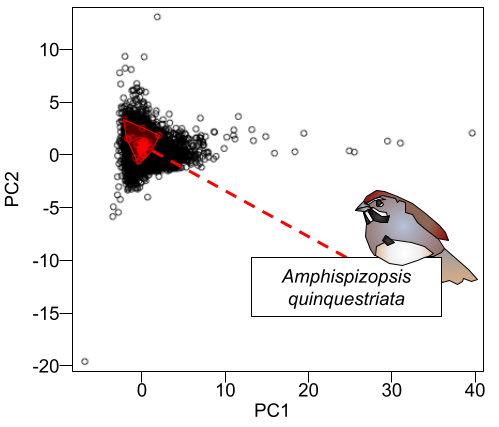

**Fig. S3.**

Breadth of syllable types across Passerellidae with one focal species highlighted. X-axis and y-axis depict PC1 and PC2, respectively, of five song variables (Table 1). Each point is an individual recording, with value showing the centroid across all syllables in that recording. Highlighted in red is one species, *Amphispizopsis quinquestriata*, with all recordings and their area given.

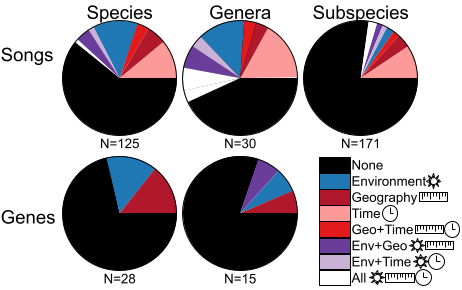

**Fig. S4.**

Proportion of species (left), genera (center), and subspecies (right) whose songs (top) or genes (bottom) are explained by different spatiotemporal factors. Slices of pie charts indicate relative proportion with colors indicating whether environment, geography, time, or a combination best explain variation. Colors indicate the significant predictors (see legend, right) -- shades of red are stochastic features (geography, time), shades of blue are environmental features, purple/white indicate some combination, and black indicates no such feature. Sample sizes are given below each plot.

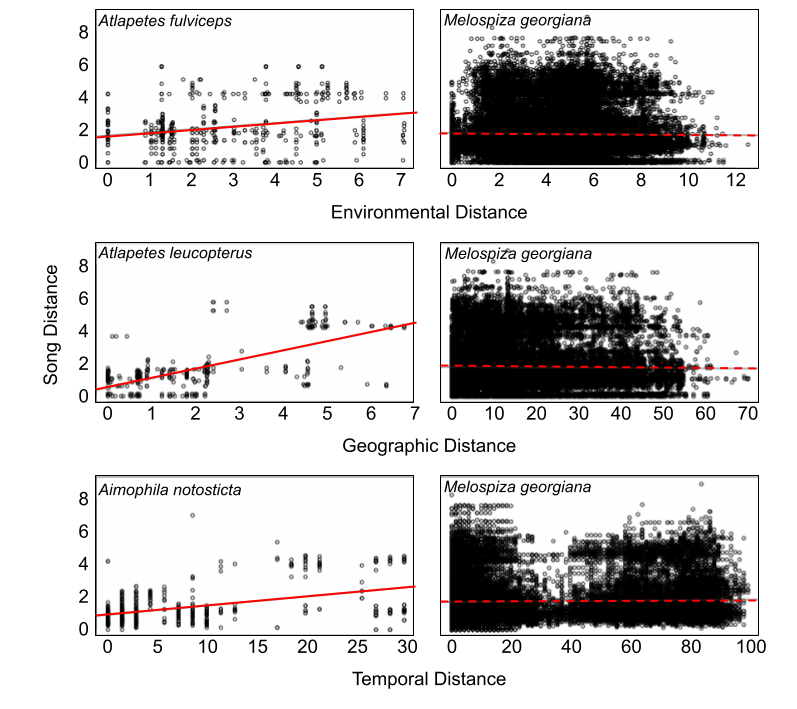

**Fig. S5.**

Relationship between song distance (y-axis) and spatiotemporal variables (x-axes) in four species. Top two panels show song vs environmental distance, middle two panels show song vs geographic distance, and bottom two panels show song vs temporal distance. Left column shows three species where the relationship is significant (solid line), the right column shows one species where relationships are always non-significant (dashed line).

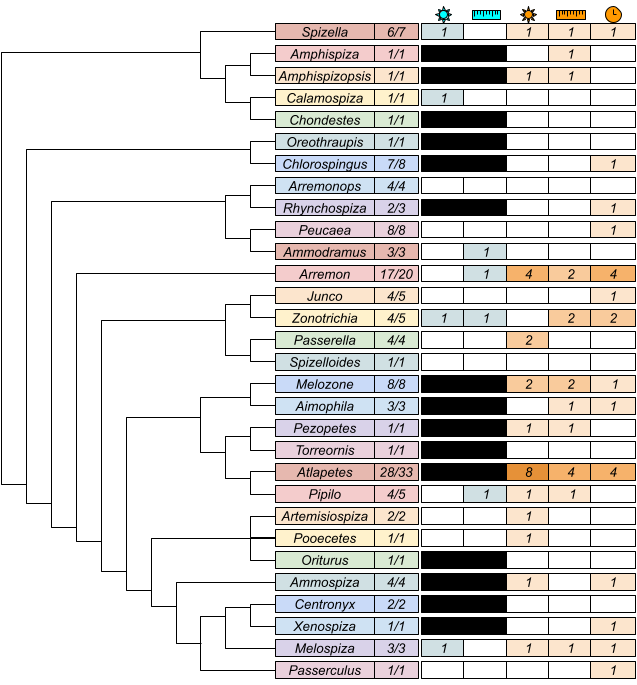

**Fig. S6.**

Best predictors of species-level differences in traits. Left: phylogeny at the generic level, with sample sizes of species sampled/total species. Right: heat map of number of species sampled that show significant differences in song (orange shades) and genetics (teal shades). Icons indicate spatial/temporal features tested: sun indicates environment, ruler indicates geography, and clock indicates time.

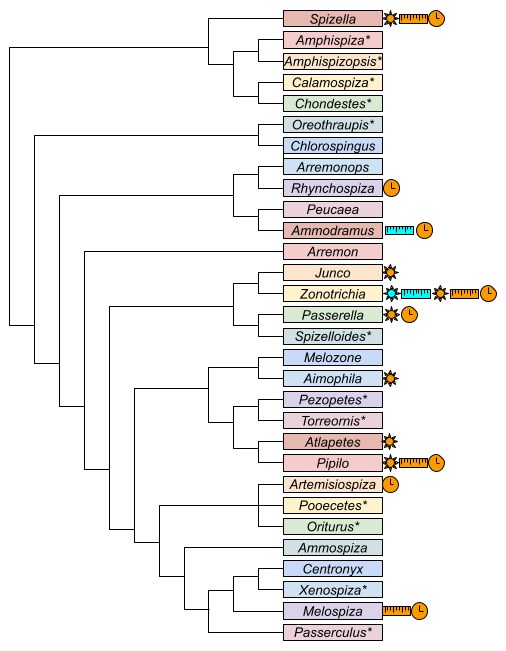

**Fig. S7.**

Best predictors of genus-level differences in traits. Icons to the right of names indicate significant associations (see Fig. S5). Stars indicate monotypic genera, so no icons are given as results are the same as in single-species results. Icons indicate spatial/temporal features tested: sun indicates environment, ruler indicates geography, and clock indicates time.

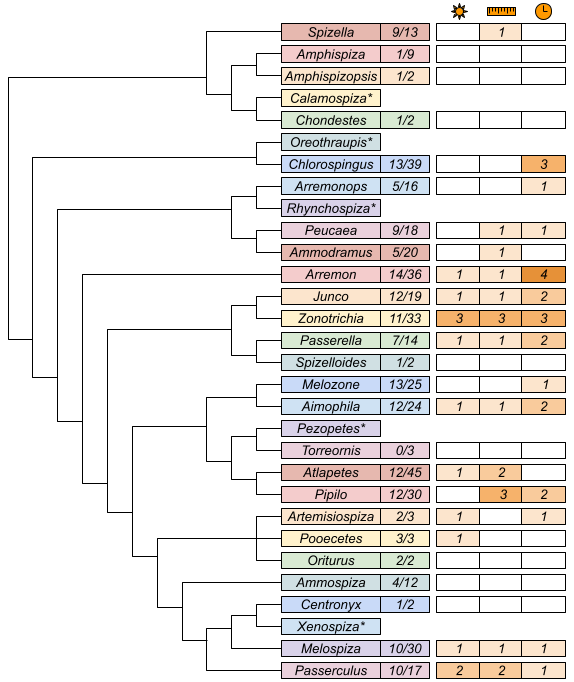

**Fig. S8.**

Best predictors of subspecies-level differences in traits. Icons as given in Fig. S5. Note that stars indicate genera with no subspecies, which are left blank. Sample sizes are given as subspecies sampled/total subspecies. Icons indicate spatial/temporal features tested: sun indicates environment, ruler indicates geography, and clock indicates time.

**Table S1.**

Principal components analysis on five song variables extracted from Passerellidae syllables. Importance of each PC and loadings/rotation of each variable given.

|  |  | **PC1** | **PC2** | **PC3** | **PC4** | **PC5** |
| --- | --- | --- | --- | --- | --- | --- |
| Importance | St. Dev. | 1.245 | 1.023 | 0.994 | 0.990 | 0.660 |
|  | Prop. Of Variance | 0.310 | 0.209 | 0.198 | 0.196 | 0.087 |
|  | Cumulative Prop. | 0.310 | 0.519 | 0.717 | 0.913 | 1.000 |
| Rotation | Bandwidth | -0.296 | 0.573 | -0.304 | -0.628 | 0.311 |
|  | Time | 0.610 | -0.001 | -0.241 | -0.465 | -0.594 |
|  | Center | 0.146 | 0.702 | 0.641 | 0.149 | -0.228 |
|  | Inflection | 0.710 | 0.023 | 0.061 | 0.004 | 0.701 |
|  | Slope | 0.121 | 0.421 | -0.659 | 0.606 | -0.083 |

**Table S2.**

Importance of first five PCs in a principal components analysis on environmental variables.

|  | **PC1** | **PC2** | **PC3** | **PC4** | **PC5** |
| --- | --- | --- | --- | --- | --- |
| Standard deviation | 3.10 | 2.39 | 2.00 | 1.93 | 1.56 |
| Proportion of Variance | 0.20 | 0.12 | 0.08 | 0.08 | 0.05 |
| Cumulative Proportion | 0.20 | 0.32 | 0.40 | 0.48 | 0.53 |

**Table S3.**

Rotation/loadings of first five PCs in a principal components analysis on environmental variables. Variables are categorized by general type.

|  |  | **PC1** | **PC2** | **PC3** | **PC4** | **PC5** |
| --- | --- | --- | --- | --- | --- | --- |
| Land Cover % | Bare | 0.076 | -0.115 | -0.060 | 0.039 | -0.110 |
|  | Built Up | 0.008 | -0.004 | 0.083 | 0.159 | 0.330 |
|  | Crops | 0.029 | -0.026 | 0.163 | -0.090 | -0.016 |
|  | Grasses | 0.116 | -0.114 | -0.052 | -0.028 | -0.273 |
|  | Moss/Lichens | 0.013 | 0.021 | -0.015 | 0.018 | -0.042 |
|  | Permanent Water | 0.029 | 0.024 | 0.059 | 0.065 | 0.006 |
|  | Seasonal Water | 0.017 | -0.011 | 0.054 | 0.007 | -0.014 |
|  | Shrubs | 0.018 | -0.231 | -0.163 | -0.028 | -0.137 |
|  | Trees | -0.136 | 0.197 | -0.050 | -0.089 | 0.015 |
| Climate | Mean Annual Temp. | -0.234 | -0.245 | 0.095 | -0.092 | 0.097 |
|  | Mean Diurnal Temp. Range | 0.137 | -0.220 | -0.134 | -0.184 | -0.071 |
|  | Isothermality | -0.264 | -0.123 | -0.103 | 0.044 | -0.123 |
|  | Temp. Seasonality | 0.276 | 0.110 | 0.097 | -0.138 | 0.039 |
|  | Max Temp. Warmest Month | 0.005 | -0.267 | 0.139 | -0.282 | 0.134 |
|  | Min Temp. Coldest Month | -0.282 | -0.176 | 0.019 | 0.037 | 0.054 |
|  | Annual Temp. Range | 0.284 | 0.046 | 0.049 | -0.174 | 0.011 |
|  | Mean Temp. Warmest Quarter | -0.106 | -0.176 | 0.196 | -0.223 | 0.046 |
|  | Mean Temp. Driest Quarter | -0.202 | -0.214 | -0.072 | 0.052 | 0.064 |
|  | Mean Temp. Warmest Quarter | -0.072 | -0.244 | 0.206 | -0.245 | 0.168 |
|  | Mean Temp. Coldest Quarter | -0.272 | -0.210 | 0.010 | 0.006 | 0.045 |
|  | Annual Precip. | -0.270 | 0.163 | 0.084 | -0.062 | -0.077 |
|  | Precip. Wettest Month | -0.282 | 0.076 | 0.018 | 0.002 | -0.058 |
|  | Precip. Driest Month | -0.136 | 0.241 | 0.185 | -0.171 | -0.079 |
|  | Precip. Seasonality | -0.060 | -0.252 | -0.170 | 0.200 | 0.079 |
|  | Precip. Wettest Quarter | -0.279 | 0.091 | 0.024 | 0.006 | -0.055 |
|  | Precip. Driest Quarter | -0.149 | 0.242 | 0.182 | -0.167 | -0.086 |
|  | Precip. Warmest Quarter | -0.204 | 0.100 | 0.124 | -0.189 | -0.141 |
|  | Precip. Coldest Quarter | -0.200 | 0.165 | 0.023 | 0.083 | -0.006 |
| Urbanization (by decade) | % Cropland | 0.056 | -0.061 | 0.242 | -0.148 | -0.040 |
|  | % Grazing Land | 0.009 | -0.207 | 0.005 | -0.074 | -0.206 |
|  | % Rice Cover | -0.001 | -0.001 | 0.009 | -0.003 | -0.005 |
|  | Census Population | -0.002 | -0.016 | 0.055 | 0.103 | 0.297 |
|  | % Land Irrigated | 0.003 | -0.117 | 0.057 | -0.026 | -0.031 |
|  | % Urban Landcover | 0.017 | -0.017 | 0.104 | 0.158 | 0.381 |
| Habitat Classifications | Biosphere-Atmosphere Transfer Scheme | -0.009 | 0.002 | 0.088 | 0.112 | -0.070 |
|  | International Geosphere Biosphere Programme | 0.018 | -0.069 | 0.363 | 0.238 | -0.140 |
|  | USGS Land Use System | -0.033 | 0.121 | -0.202 | 0.031 | -0.143 |
|  | Olson Global Ecosystem | -0.001 | -0.133 | 0.055 | -0.111 | -0.218 |
|  | Simple Biosphere 2 Model 2 | 0.072 | -0.106 | 0.287 | 0.268 | -0.234 |
|  | Simple Biosphere Model 1 | 0.052 | -0.068 | 0.304 | 0.297 | -0.224 |
|  | Running Vegetation Lifeforms | 0.017 | -0.050 | 0.269 | 0.225 | -0.219 |
| Topography | Elevation | -0.003 | -0.036 | -0.267 | -0.038 | -0.311 |
|  | Relief | -0.031 | 0.006 | -0.013 | 0.024 | -0.044 |
|  | Roughness | -0.071 | 0.018 | -0.002 | 0.357 | 0.120 |
|  | Ruggedness Index | -0.149 | -0.031 | -0.214 | 0.129 | -0.074 |
|  | Slope | -0.063 | -0.022 | -0.158 | 0.103 | -0.005 |
| Soil | Acidity | -0.165 | 0.244 | 0.026 | -0.050 | -0.015 |
|  | Texture Classification | 0.143 | 0.154 | -0.008 | 0.068 | 0.027 |
